# Genetic ablation of a female specific Apetala 2 transcription factor blocks oocyst shedding in *Cryptosporidium parvum*

**DOI:** 10.1101/2022.11.23.517783

**Authors:** Jayesh Tandel, Katelyn A. Walzer, Jessica H. Byerly, Brittain Pinkston, Daniel P. Beiting, Boris Striepen

**Affiliations:** Department of Pathobiology, School of Veterinary Medicine, University of Pennsylvania, Philadelphia, PA; Center for Tropical and Emerging Global Diseases, University of Georgia, Athens, GA

## Abstract

The apicomplexan parasite *Cryptosporidium* is a leading global cause of diarrheal disease, and the infection poses a particularly grave threat to young children and those with weakened immune function. Infection occurs by ingestion of meiotic spores called oocysts, and transmission relies on fecal shedding of new oocysts. The entire lifecycle thus occurs in a single host and features asexual as well as sexual forms of replication. Here we identify and locus tag two Apetala 2-type (AP2) transcription factors and demonstrate that they are exclusively expressed in male and female gametes, respectively. To enable functional studies of essential genes in *C. parvum* we develop and validate a small molecule inducible gene excision system, which we apply to the female factor AP2-F to achieve conditional gene knock out. Analyzing this mutant, we find the factor to be dispensable for asexual growth and early female fate determination in vitro, but to be required for oocyst shedding in infected animals in vivo.

Transcriptional analyses conducted in the presence or absence of AP2-F revealed that the factor controls the transcription of genes encoding crystalloid body proteins, which are exclusively expressed in female gametes. In *C. parvum*, the organelle is restricted to sporozoites, and its loss in other apicomplexan parasites leads to blocked transmission. Overall, our development of conditional gene ablation in *C. parvum* provides a robust method for genetic analysis in this parasite that enabled us to identify AP2-F as an essential regulator of transcription required for oocyst shedding and transmission.

## INTRODUCTION

Parasites rely on complex lifecycles to colonize a variety of hosts, organs, and tissues, to massively amplify their numbers in the course of infection, and to enable transmission between hosts. The lifecycles are defined by developmental progression through a series of distinct morphological stages, each highly adapted to specific environments and tasks. Examples include the formation of chronic tissue cysts for spread via carnivory, or aggregation in the salivary glands of blood feeding insects to enable vector-borne transmission. For members of the protist phylum Apicomplexa, these lifecycles also alternate phases of clonal amplification through asexual cell division, and sexual processes characterized by differentiated male and female gametes, fertilization, genomic recombination, and meiosis (1). Often, lifecycle completion requires transmission between multiple hosts. In the case of *Plasmodium falciparum*, the causative agent of the most dangerous form of malaria, infection of humans and *Anopheles* mosquitoes is required to complete the life cycle. In contrast, *Cryptosporidium* has a much simpler single host lifecycle where both asexual and sexual stages are found in the same host and the same tissue (2).

*Cryptosporidium* is one of the most important causes of severe diarrheal disease under a variety of epidemiological settings around the world (3-5). Cryptosporidiosis is life threatening in immunosuppressed individuals, like those with HIV-AIDS or transplant recipients. Immunocompetent individuals are susceptible, but the infection is self-limiting. In the United States, *Cryptosporidium* is responsible for 50% of disease outbreaks linked to recreational water use, and is overall the most important cause of waterborne illness (5-7). Effective treatment is lacking, and more potent drugs are urgently needed (3, 4). An important lesson for *Cryptosporidium* drug development from recent advances in the search for anti-malarials is to broaden the attack over the entire lifecycle (9). In *Cryptosporidium* this is of even greater importance as sustained infection is believed to be fed by successive cycles of autoinfection (8, 9). Currently, there is no vaccine to prevent cryptosporidiosis, but there is evidence that children develop protective natural immunity (10). The sexual phase of the lifecycle offers attractive targets for vaccination, and both vaccine and drug development depend on a better understanding of the complete lifecycle.

*Cryptosporidium* oocysts ingested with water or food release motile sporozoites into the intestinal tract which proceed to invade enterocytes, where they occupy an intracellular but extra-cytoplasmic apical niche (11, 12). Asexual replication (merogony) results in release of a new generation of motile and invasive forms leading to the rapid spread of the parasite through the intestine. An intrinsic program of developmental commitment then leads to differentiation into male (micro) and female (macro) gametes (13). Male gametes egress to fertilize female gametes, which remain in their host cell. The resulting intracellular zygote undergoes meiosis and develops into a new oocyst. Once released from its host cell, the oocyst is either shed with the feces, resulting in transmission, or sporozoites emerge to reinitiate the cycle within the same host. Oocysts with distinct wall structure have been morphologically described (8) and this has been suggested to reflect adaptation to differential transmission fates.

In the human cell line HCT-8, *C. parvum* initially vigorously amplifies through three rounds of asexual replication. After 48 hours, parasites differentiate into gametes in a synchronous and highly predictable fashion (2, 13-15) and they do so directly from 8N type-one meronts. However, while gametes form and meet in HCT-8 culture, fertilization fails, no new oocysts are formed, and parasites cease to proliferate (2). Longer term proliferation has been achieved in stem cell derived culture models, coinciding with sex and oocyst formation (16-18). This may suggest that *Cryptosporidium* sex contributes to continued infection of a single host in addition to its well-established role in transmission between hosts.

Progression through the apicomplexan lifecycle is accompanied by dramatic changes in gene expression that often are the consequence of regulation at the level of transcriptional initiation (19); however, there are also examples of control through translational repression (20, 21). Initiation is governed by promoter elements immediately 5’ of apicomplexan genes that serve as landing spots for the transcription machinery and its regulatory components, and sequence specific interaction between promoters and different transcription factors enable stage-specific expression patterns. Apetala-2-type (AP2) DNA binding proteins (22), in particular, have emerged as transcriptional modulators with central roles in apicomplexan lifecycle progression. They have been linked to developmental regulation of gametocytes (23, 24), liver stages (25), ookinetes (26), and sporozoites (27) in *Plasmodium spp*., and tissue cyst formation in *Toxoplasma gondii* (28). The *C. parvum* genome encodes 16 putative ApiAP2 proteins (29, 30), some of which appear transcribed in a stage-specific fashion (2).

Here we report the identification and analysis of two AP2 proteins that are exclusively expressed in male or female gametes, respectively. We found cgd4_1110, which encodes the female specific AP2-F, to be refractory to targeted disruption of the locus. To enable genetic studies of essential genes, we established small molecule inducible Cre recombinase mediated gene ablation. Using this model, we engineered mutant parasites in which the expression of cgd4_1110 is conditional and can be controlled with a small molecule. We tested the impact of loss of the protein on parasite development and overall infection *in vitro* and *in vivo*.

## RESULTS

### The AP2 DNA binding protein cgd4_1110 is exclusively expressed in female gametes

Little is known about the transcriptional control of lifecycle progression in *Cryptosporidium* and the factors that orchestrate the commitment to, and the execution of, different developmental cell fates. However, the *C. parvum* genome encodes numerous AP2 DNA binding proteins, homologs of which have been linked to stage-specific gene expression control in the related parasite *Plasmodium* (19). Analysis and experimental manipulation of these factors thus may provide insight into the *Cryptosporidium* life cycle. Initial PCR screening at different times of culture suggested that some of these factors themselves may underlie transcriptional control across the lifecycle (29, 30). We recently used engineered reporter parasites to isolate specific stages of *Cryptosporidium* and carried out comparative transcriptomics by RNA-seq (2). This analysis suggested five AP2s to be preferentially transcribed in sexual stages (cgd1_3520, cgd2_3490, cgd4_1110, cgd6_2670, and cgd8_810). We initially focused on cgd4_1110, which appeared to be transcribed in female gametes, and tagged the gene to yield C-terminal translational fusion with either an mNeon-Green reporter, or a triple HA epitope tag introduced into the native locus by Cas9 mediated homologous recombination (31) (see Supplementary Fig. 1 for maps and PCR validation). cgd4_1110-3xHA parasites were used to infect HCT-8 cultures, which were subsequently fixed at 24 and 48h post-infection, stained with an HA-specific antibody, and counter-stained with *Vicia villosa* lectin, which recognizes glycan epitopes found on all *C. parvum* stages. No HA staining was observed at 24h, a timepoint at which only asexual forms are found. However, at 48h, when differentiation to gametes has occurred, 60.7% (+/-4,4%) of parasites were HA positive (Fig. 1 A and B). Using cgd4_1110-mNG parasites, we similarly found label only at the 48h-timepoint. Relying on nuclear morphology to distinguish stages (2, 13) at this timepoint, we detected the reporter in female gametes, but not in asexual meronts or in male gamonts (Fig. 1C). Consistent with the presumptive function of a DNA-binding protein, the label coincided with the single nucleus of the female gamete (arrowhead Fig. 1C). To further validate this finding using a molecular marker, we infected cells with cgd4_1110-HA parasites and stained with antibodies to HA and the meiotic recombinase DMC1, which was recently shown to be restricted to female gametes (15). We observed HA staining exclusively in DMC1 positive cells (Fig. 1D).

**Figure 1:**
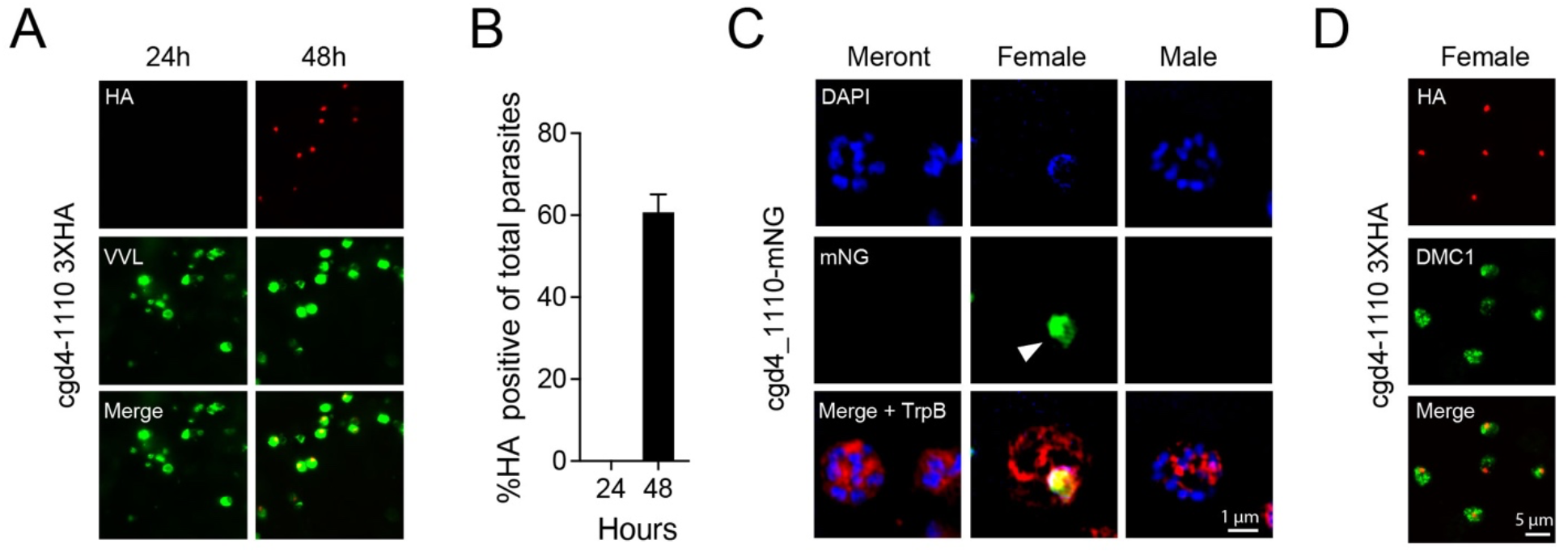
*C. parvum* cgd4-1110 encodes an AP2 DNA binding protein that is expressed exclusively in female gametes. (**A**) The locus of cgd4_1110 was modified to append a 3xHA epitope tag to the coding sequence. The resulting parasite strain was used to infect HCT-8 cell cultures and analyzed by immunofluorescence assays (IFA) at 24 and 48h post infection. The transgene was detected with antibodies to HA (red) and all parasites were labeled with *Vicia villosa* lectin (VVL, green). (**B**) Quantification of HA staining as a proportion of all parasites. (**C**) IFA of cultures infected with cgd4_1110-mNG showing asexual meronts, a female gamete, and a male gamont at 48h. mNeon Green (mNG) fluorescent protein is shown in green, DNA stained with DAPI in blue, and TrpB (labeling all parasites) in red. The single nucleus of the female gamete is highlighted with an arrowhead. (**D**) cgd4_1110-3xHA (red) expressing parasites stained for the female marker DMC1 (note that all HA positive parasites are also DMC1 positive).

### Male gamonts transiently express the AP2 protein cgd6_2670

Previous transcriptional profiling work by us and others suggested that, due to timing of expression, cgd6_2670 might also be a female specific AP2 (2, 30). To further test this, the locus was tagged with a triple HA epitope or an mCherry reporter, and cultures infected with the resulting parasite strains were fixed at different timepoints. Upon staining, we observed fluorescence exclusively at the 48 h timepoint as expected (Fig. 2A); however, compared to cgd4_1110, this staining was limited to a much smaller proportion of the overall parasite population (Fig. 2B). Analysis of the mCherry strain showed nuclear fluorescence exclusively in parasites with 4 or 8 round nuclei, which appeared clustered into a rosette at the center of intracellular parasite stages (Fig. 2C). No labeling was associated with identifiable asexual stages or with female gametes, but faint cytoplasmic staining was notable in mature male gamonts. The temporal pattern of cgd6_2670 transcription matches that of the male fusion protein HAP2 (Fig. 2D). We therefore studied parasites engineered to express an HA tagged version of HAP2 (2, 32). HAP2 is found as a characteristic dot at the apex of mature male gametes, and is perinuclear in immature gamonts, likely reflecting protein in the endoplasmic reticulum *en route* through the secretory pathway (Fig. 2E). Importantly, these immature stages showed centrally rosetted nuclei. We quantified the nuclear area of these stages (1.46 ± 0.32 μm^2^) and found it similar to that of cgd6_2670 positive rosette stages (1.829 ± 0.33 μm^2^), but significantly different from that of asexual meronts (Fig. 2F; 4.12 ± 0.83 μm^2^; ^****^p< 0.0001). The nuclear morphology of these stages is reminiscent of developing male gamonts recently observed by time lapse imaging (15). Next, we engineered parasites to express a tdNeon reporter under the control of the cgd6_2670 promoter to mark all stages in which this gene is transcribed. In contrast to our previous experiments, we did not fuse the reporter to the AP2 protein itself, thus releasing it from posttranslational control through development (see Fig. 2I). We introduced this construct into the HAP2 locus, adding an HA tag to specifically identify males. When studying the resulting transgenic strain, we again found fluorescence to be restricted to the 48h time point, but now observable in higher numbers, matching those previously observed for male gamonts (Fig. 2G-J (2)). Importantly, essentially all parasites expressing tdNeon were also positive for the male marker HAP2-HA (Fig. 2K and L). We conclude that the two AP2 proteins characterized here are expressed in a female (cgd4_1110) and early-male-specific (cgd6_2670) fashion and thus named them AP2-F and AP2-M, respectively.

**Figure 2:**
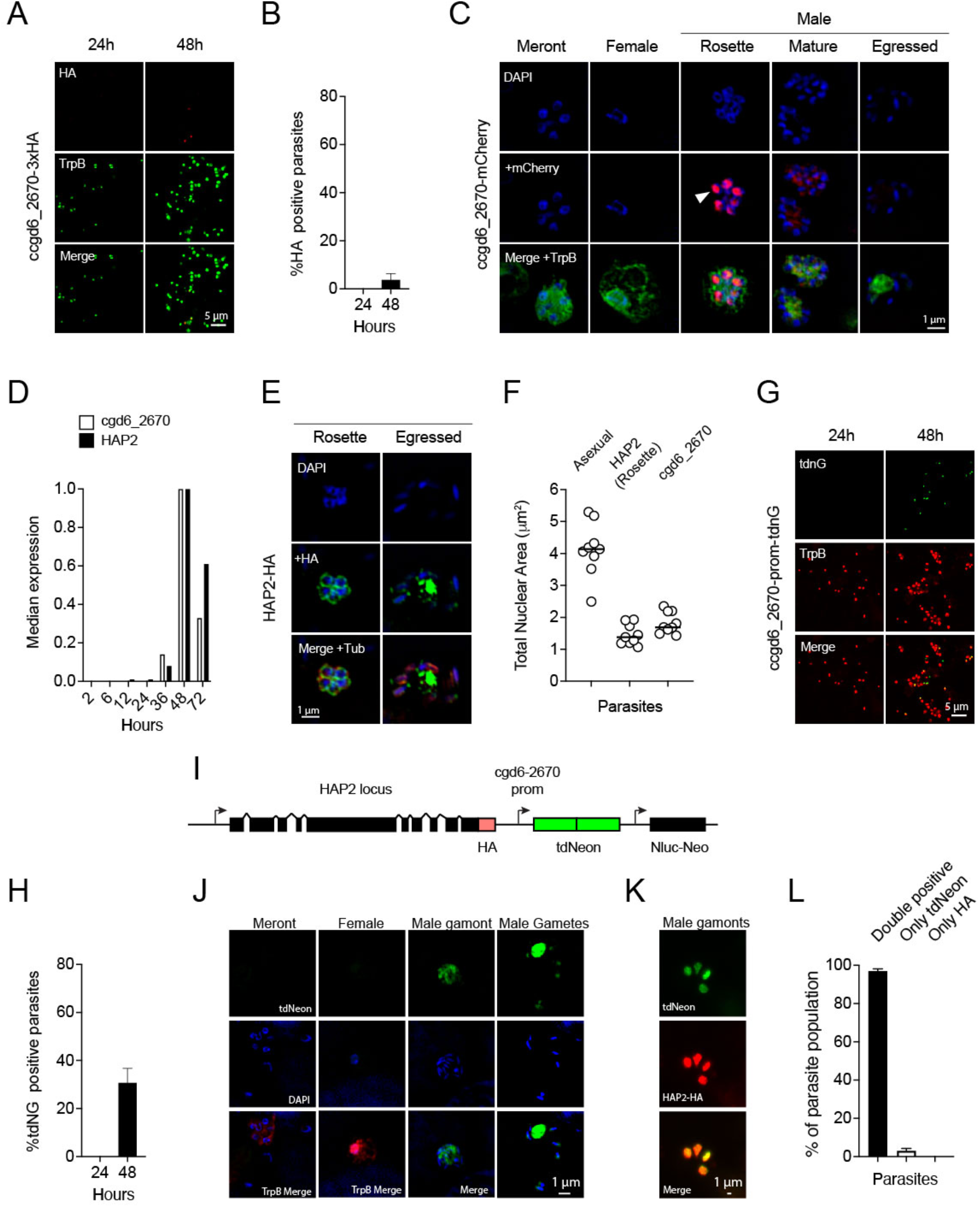
*C. parvum* cgd6_2670 encodes a cell cycle restricted male gamont specific AP2 protein. (**A**) A 3x HA tag was introduced into the cgd6_2670 locus and the resulting parasites were subjected to IFA after growth in HCT-8 cultures for the indicated time. (**B**) Quantification of HA positive parasites as a fraction of all parasites (TrpB). (**C**) Super resolution images showing parasites expressing a cgd6_2670-mCherry fusion protein (red) counter stained with TrpB (green). Note strong nuclear staining of parasite nuclear rosettes and weak labeling of mature male gamonts. (**D**) mRNA abundance for cgd6_2670 and HAP2 (qPCR data from (32)). (**E**) IFA of HAP2-HA expressing parasites showing labeled parasites with nuclear rosettes. (**F**) Quantification of nuclear area (based on DAPI staining in IFAs) for asexual meronts and nuclear rosette stages stained for cgd_2670 or HAP2. (**G**) Expression of the tdNeon fluorescent reporter under the control of the cgd6_2670 promoter at the indicated times. (**I**) Schematic view of the HAP2 locus of the strain engineered for (**G**-**K**). Note that this parasite simultaneously reports on the expression of HAP2 and cgd6_2670 but is released from posttranslational control of cgd6_2670. (**H**) Quantification of mNeon positivity over time. (**J**) IFA analysis of different stages. Note green fluorescence in immature and mature male gametes. (**K**) IFA of four male gamonts (note different scale) labeled with tdNeon and HA antibody for HAP2, (**L**) is showing quantification (n=3, bars show mean and STD).

### Conditional gene ablation in *Cryptosporidium parvum*

To understand the role of gamete specific AP2 factors in *Cryptosporidium* lifecycle progression, we next set out to disrupt their loci. An attempt to disrupt AP2-M was unsuccessful (data not shown). We next attempted to insert a drug marker into the N-terminal AP2-F DNA binding domain. This was unsuccessful in five attempts using three different guide RNAs. Parallel control experiments inserting an epitope tag without altering the coding sequence readily resulted in transgenic parasites (we note that we were able to disrupt the C-terminal DNA binding domain, this did not impact parasite growth or development and we thus did not study this mutant any further, see supplementary Fig. 2). In *C. parvum* transgenesis, selection occurs in animals, and recovery of modified parasites requires the formation of oocysts, following the completion of the full lifecycle (31). While failure to disrupt the locus is consistent with essentiality of a gene, it is not proof, and we thus next sought to develop an experimental model to ablate genes in *C. parvum* conditionally. Multiple approaches have been adapted to apicomplexans for this purpose, including modulating transcription initiation (33), affecting the stability of the targeted mRNA or protein (34-37), or using recombinases to excise portions of the genome (38, 39).

Here we used a modified version of Cre-recombinase (DiCre) that, upon rapamycin induction, assembles an active enzyme from two fragments to excise gene segments flanked by LoxP recognition sequences (38, 40). To enable the introduction of loxP sites into the coding sequence of genes without changing their protein products, we explored the use of introns. We searched the *C. parvum* genome for short introns and through preliminary experimentation settled on the 73 bp intron from cgd1_1320, which is expressed across the life cycle (32). To test the functionality of this intron in trans, we inserted it into the nanoluciferase (Nluc) coding region (Fig. 3A). Transient transfection of this reporter into sporozoites followed by infection of HCT-8 cells resulted in luciferase activity indistinguishable of that obtained with a continuous Nluc coding sequence (Fig. 3B). Faithful splicing was critical for this, as the activity was lost when the 5’ splice donor site of the intron was ablated (Fig. 3B; *p*< 0.0001). Next, loxP sequences were introduced at different positions of the intron and luciferase activity was measured. We identified multiple positions that resulted in comparable activity to that of the native intron (Fig. 3C, see supplementary Fig. 3 for detail). We conclude that the intron is faithfully spliced when introduced in trans, and its splicing remains intact when loxP sites are incorporated.

**Figure 3:**
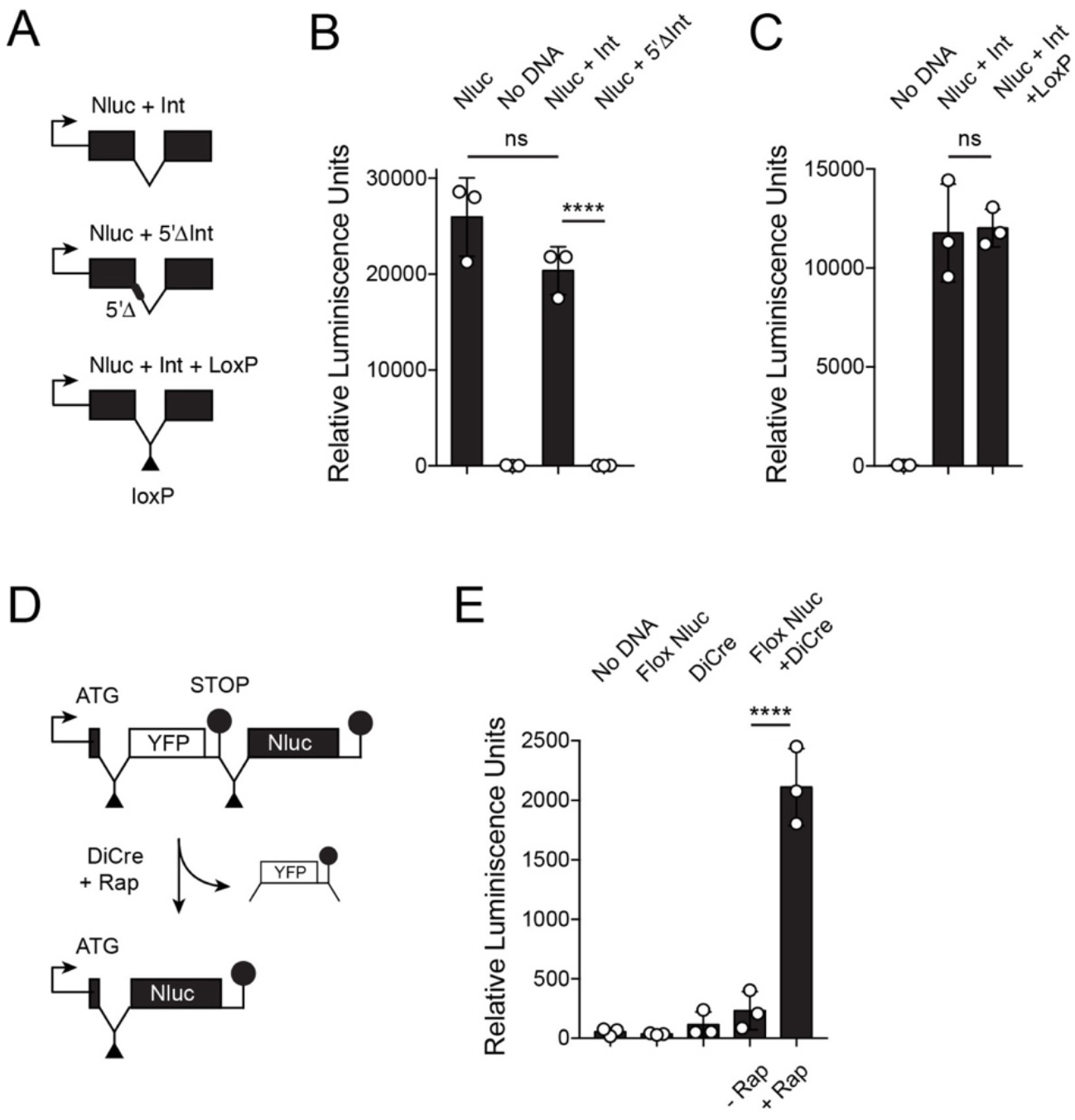
Cre mediated DNA excision using a floxed intron sequence. (**A**) Schematic maps of Nluc reporters used in transient transfection experiments. (**B** and **C)** Sporozoites were transfected with the indicated constructs (see Materials and Methods for detail) and used to infect HCT-8 cell cultures. Luciferase activity was measured after 48h of culture. Note that introns and loxP sites within introns do not perturb Nluc expression. (**D**) Schematic outline of the transient Cre recombinase assay. Rapamycin (Rap) induction of Di-Cre results in excision of a floxed stop sequence (YFP) and expression of Nluc. (**E**) Transient transfection assays using the indicated constructs followed by luciferase measurement. Note induction of Nluc activity upon addition of rapamycin (+Rap) when DiCre and floxed reporter (Flox Nluc) plasmids were cotransfected. (**B, C**, and **E**) Each symbol represents one independent well and the bar shows the mean and standard deviation. Significance was evaluated using an unpaired Student’s t-test.

Next, we constructed a plasmid that expresses two fragments of Cre recombinase under control of the *C. parvum* enolase promoter, each fragment fused to FKBP12 or FRB, respectively, and separated by a viral translational skip sequence (40). To test whether this system results in rapamycin inducible Cre activity in *C. parvum*, we designed a transient reporter assay. In our test construct, Nluc expression was disrupted by a floxed stop sequence containing a YFP reporter (Fig. 3D). Cre-mediated excision should remove the stop sequence and result in Nluc expression. Transfection of this floxed reporter, or DiCre alone, did not result in luciferase activity above a background control. However, when parasites were simultaneously transfected with both constructs and grown in culture medium containing 100 nM rapamycin, robust activity was detected, and this activity depended on the presence of rapamycin (Fig. 3E *p*< 0.0001). Please note that rapamycin does not impede *Cryptosporidium* growth at the concentration used here. We tested a range of concentrations and determined the IC_50_ to be 8.3 μm (see Supplementary Fig. 4).

### Rapamycin-inducible gene excision from the *Cryptosporidium* genome

Next, we tested the ability of the system to excise floxed sequences from the parasite genome. We targeted the well-characterized non-essential thymidine kinase (TK) locus (41, 42), replacing the last 222 bp of the native sequence with a recodonized version preceded by an artificial intron carrying a loxP site (Fig. 4A). We also appended a 3XHA epitope tag to be able to detect the protein along with cassettes for expression of DiCre and the Nluc-Neo selection marker. PCR mapping confirmed successful modification of the TK gene (Supplementary Fig. 1), and Sanger sequencing of the amplified insert demonstrated that the targeted region was flanked by loxP sites. Using primers flanking the floxed region we detected two amplicons representing the floxed locus, prior to and following Cre mediated excision (note that all primers used in this study are listed in supplementary table 1). The initially faint smaller 743 bp band became more noticeable upon continuous passage of the strain, suggesting that stable expression of DiCre recombinase in this strain resulted in some ‘leaky’ activity in the absence of rapamycin.

**Figure 4:**
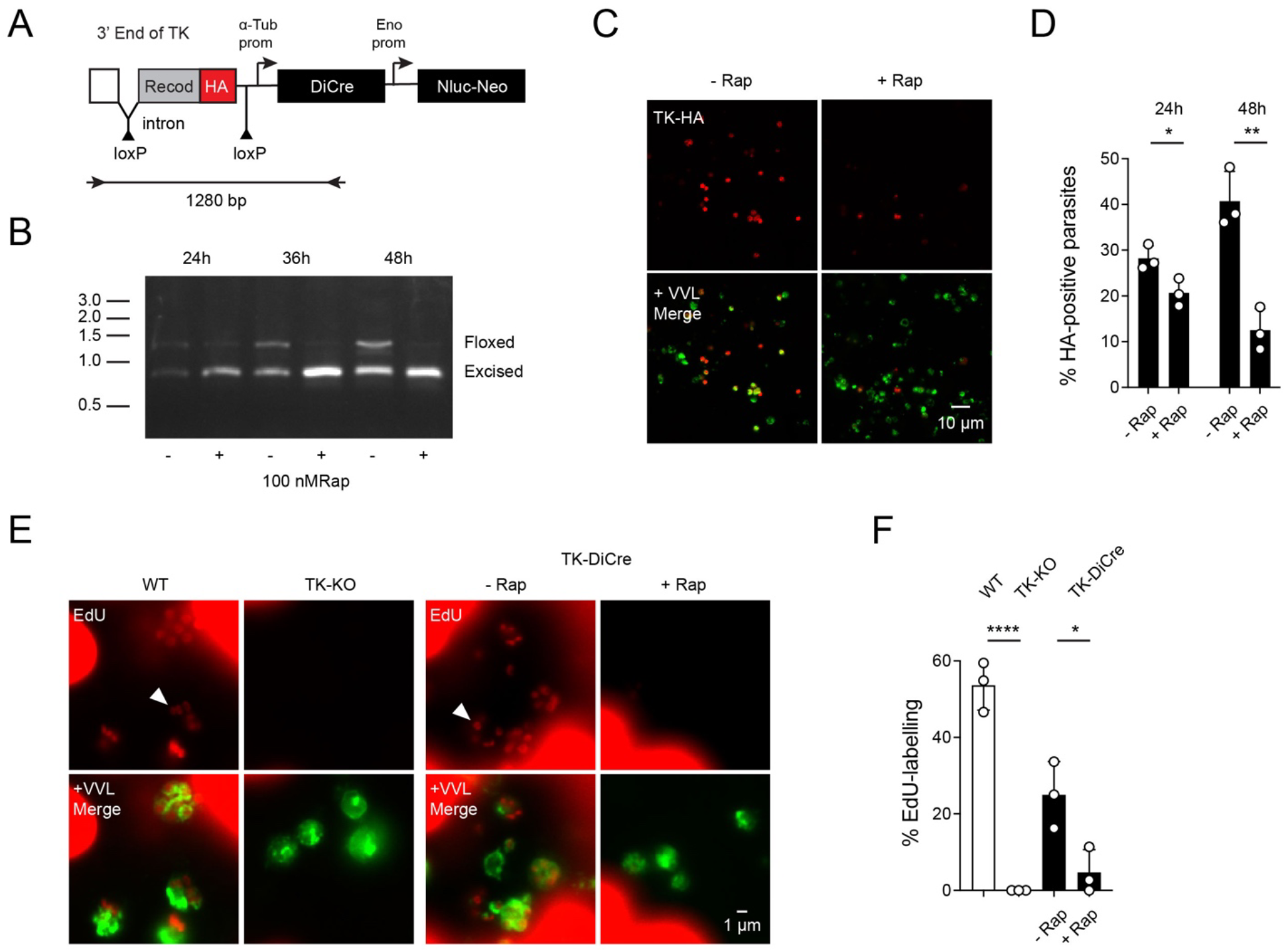
Inducible excision of a gene from the *C. parvum* genome. (**A**) Schematic view of the targeting construct used to modify the thymidine kinase (TK) locus. Note that HA tag is located between loxP sites and will be excised due to Cre activity. (**B**) Modified parasites were used to infect HCT-8 cell cultures for the indicated times in the presence or absence of rapamycin. Genomic DNA was extracted and used for PCR analysis. Diagnostic amplicons for the floxed and excised locus are highlighted. (**C**) Infected cultures grown in the presence or absence of rapamycin for 48h prior to IFA and staining with antibody to HA (red) and VVL (green). (**D**) Quantification of IFA experiments showing HA positive cells as a fraction of all parasites (labeled with VVL). (**E**) 5-ethynyl-2′-deoxyuridine (Edu, red) incorporation assay, which requires TK activity, parasite nuclei are highlighted by arrowheads. Parasites were counterstained with VVL (note that the much larger host cells robustly incorporate Edu as well, this remains unchanged by rapamycin treatment). (**F**) Quantification of Edu incorporation comparing wild type (WT), a mutant in which the entire locus was replaced with a marker gene (TK-KO), and the inducible mutant (TK-DiCre) grown in the presence or absence of rapamycin. Each symbol represents an independent culture, the bar shows the mean with standard deviation. Significance was evaluated using unpaired Student’s t-test.

We next infected HCT-8 cells with TK DiCre parasites and cultures were harvested after 24, 36, and 48 hours +/- rapamycin, and genomic DNA was isolated and analyzed by PCR for the presence of the floxed region (Fig. 4B). Rapamycin treatment resulted in progressive loss of the full-length band beginning at 24 hpi. We next scored parasites in immunofluorescence experiments using anti-HA antibody to detect the (floxed) epitope and *Vicia villosa* lectin (VVL) as counterstain for all parasites. As expected from the PCR mapping, we detect both tagged and untagged parasites in cultures grown in the absence of rapamycin. When rapamycin was added to cultures, we detected significant loss of HA staining (Fig. 4C and D; Control v/s Rapamycin= 12.53% ± 4.67% after 48h of rapamycin compared to 40.72% ± 6.5% in controls; *p*< 0.01). We also directly measured the biochemical activity of TK using EdU incorporation. EdU incorporation relies on TK to activate the probe to nucleotide level (31, 43). WT parasites and a mutant lacking TK (31) were used as positive and negative controls, respectively. 53.64% ± 6.5% of the WT parasites were EdU-positive, while no staining was observed in the TK KO (Fig. 4E and F). In the TK DiCre strain 24.98% ± 8.7% of the untreated parasites were found to be EdU-positive, and this number dropped to 4.7% ± 5.94% upon rapamycin treatment (Fig. 4F, *p*< 0.05). We conclude that *C. parvum* genes tolerate a floxed intron retaining normal transcription and translation, and that treatment with rapamycin induces loss of the targeted gene, and its encoded protein and activity. We also noted some leaky DiCre activity when targeting a non-essential gene.

### Inducible loss of AP2-F through promoter ablation

We next sought to impose rapamycin inducible ablation on the likely essential AP2-F gene. Our initial attempts to flox the entire gene while concurrently introducing cassettes for DiCre and selection marker failed. In our experience, targeting cassettes exceeding 5 kb in size often fail to recombine efficiently. We thus decided to flox the promoter and initiation codon, a much shorter, but nonetheless essential sequence (see schematic in Fig. 5A). To avoid undue recombination, we typically recodonize homologous sequences. This is not feasible for a promoter, and we therefore replaced the native AP2F promoter with a floxed promoter with matching transcriptional profile. We chose the promoter of the *Cryptosporidium* oocyst wall protein 1 (COWP1), which is active exclusively in female gametes (2), and we also added an N-terminal 3XHA epitope tag. Transfection with this construct produced viable drug-resistant parasites, and PCR mapping and Sanger sequencing confirmed the modification, indicating that the promoter swap was tolerated (Supplementary Fig. 1). HCT-8 cells were infected with the AP2-F floxed strain and incubated for 24, 36, and 48 hours in the presence or absence of 100 nM rapamycin. PCR analysis of genomic DNA isolated from these cultures, using primers spanning both loxP sites, found a single band in parasites grown in the absence of rapamycin (Fig. 5B). Growth in presence of rapamycin resulted in the appearance of a second smaller band, indicating promoter excision. To assess the impact of rapamycin treatment on the level of AP2-F protein, infected HCT-8 cultures were stained with antibodies to HA (AP2-F) and DNA meiotic recombinase 1 (DMC1, a marker for female gametes (15), Fig. 5C). Rapamycin treatment resulted in significant loss of AP2-F staining, 7.62 ± 3.38% compared to 60.22 ± 18% in untreated controls (Fig. 5D, *p*< 0.01), but did not impact on parasite growth in culture (Fig. 5G). As shown in Fig. 5E and F loss of AP2-F did not result in loss of female parasites, suggesting that AP2-F does not act as a female commitment factor.

**Figure 5:**
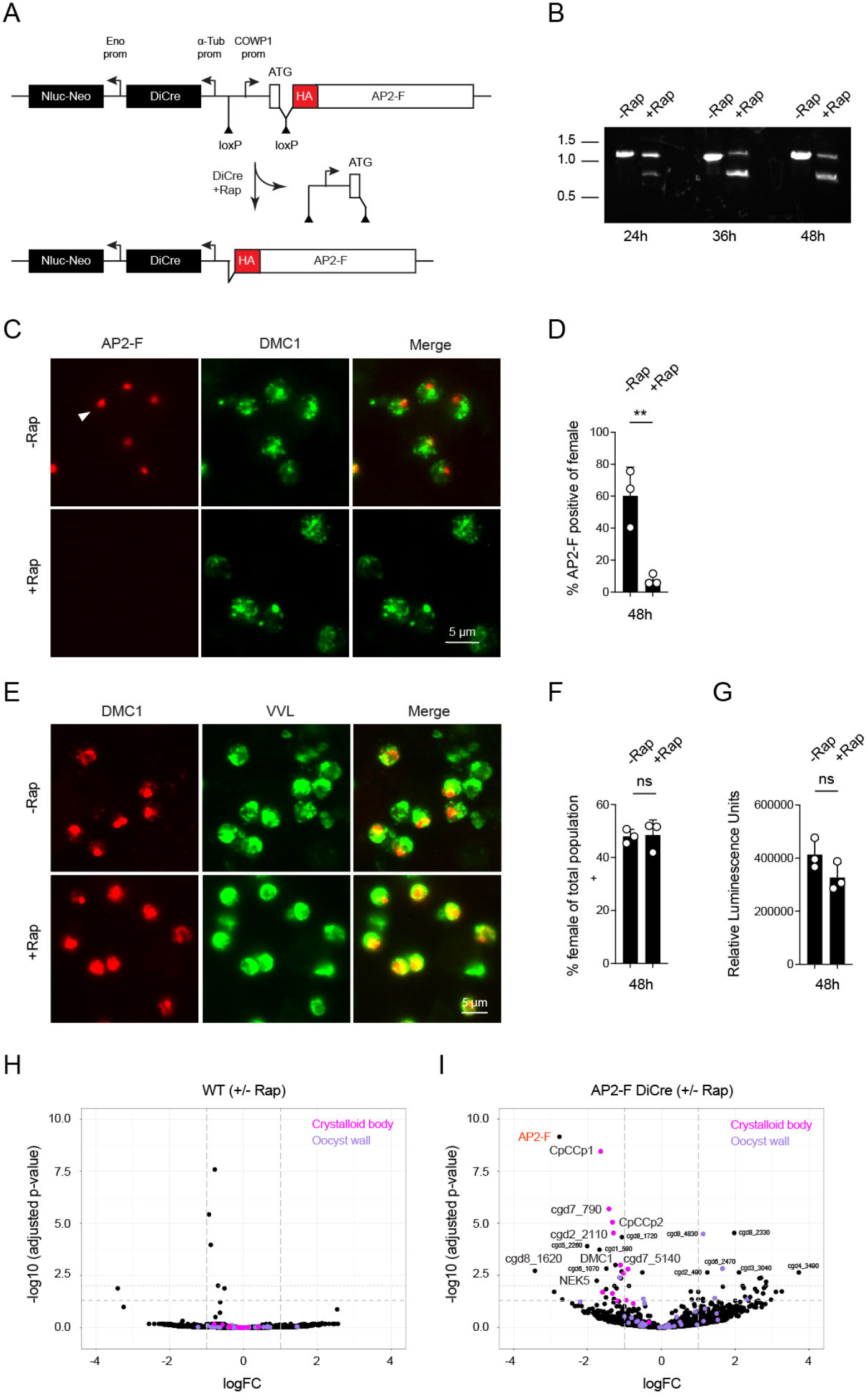
Conditional ablation of AP2-F expression by promoter excision. (**A**) Schematic view of the modified AP2-F locus prior to and after Cre mediated excision. Note that the 3’ loxP site was introduced into the coding sequence within an artificial intron and that excision removes both promoter and initiation codon. (**B**) Modified parasites were used to infect HCT-8 cell cultures for the indicated times in the presence or absence of rapamycin. Genomic DNA was extracted and used for PCR analysis. Diagnostic amplicons are highlighted. (**C**) IFA after 48h with or with rapamycin. AP2-F is labeled by HA (red, note staining of single nuclei) and female parasites were labeled with an antibody to DMC1 (green). (**D**) Quantification of AP2-F positive parasites as a fraction of all female parasites at 48h. (**E** and **F**) IFA scoring the proportion of all AF2-F/DiCre parasites (VVL, green) that develop into female gametes (DMC1, red) after 48h when grown with or without rapamycin. (**G**) Luciferase assay measuring growth of AP2-F/DiCre parasites after 48h with or without rapamycin. Each symbol represents an independent culture and the bar shows the mean with standard deviation. Significance was evaluated using an unpaired student’s t-test. WT (**H**) or AP2-F/DiCre **(I**) parasites were used to infect HCT-8 cultures. After 48h of growth, in the presence or absence of rapamycin, RNA was extracted and sequenced. Volcano plots show the relative abundance of transcript with and without treatment. Note lack of change for the WT and enrichment of crystalloid body components versus oocyst wall components (highlighted in pink and purple, respectively).

To test whether AP2-F affects gene expression and to identify genes modulated by the factor, we next performed mRNA sequencing experiments. Three biological replicates of WT and AP2-F-DiCre parasites were grown in HCT-8 cells with or without rapamycin for 48 hours, RNA was extracted, and cDNA libraries were prepared for Illumina sequencing. Comparison of WT parasites treated with and without rapamycin yielded little difference in parasite gene expression (Fig. 5H), indicating that rapamycin alone did not induce a significant transcriptional response. Upon addition of rapamycin to the AP2-F-DiCre strain, we noted reduction of transcripts for AP2-F itself by greater than 2.5-fold (Fig. 5I, highlighted in red). 15 additional genes showed significantly decreased expression and 17 genes increased in RNA abundance (see Fig. 5I and supplementary file 1). 14 of the 16 genes with reduced expression upon treatment showed tight specificity for female gametes in a previous study (2), while only six of 17 with increased expression were restricted to female gametes. Most prominent among the female specific transcripts was a cluster of crystalloid body proteins (all proteins linked to this sporozoite-specific organelle in a recent proteomic study (44) are highlighted in magenta in Fig. 5H and I). Two proteins involved in meiosis, DMC1 and NIMA kinase 5, also showed reduction as did a DNA polymerase. We note that other abundant classes of female proteins, like those that make up the oocyst wall proteome (highlighted in purple (44)), remained unchanged, suggesting specificity for a subset of female genes.

### Loss of AP2-F results in a loss of oocyst shedding in vivo

Our initial KO experiments suggested AP2-F to be an essential gene, but while we noted changes in female transcription in culture upon rapamycin treatment, these changes did not result in obvious changes in either growth or differentiation over the development observable in HCT-8 cell culture (fertilization, meiosis, and oocyst formation and release do not occur in this system (2)). We thus considered that the processes governed by AP2-F, including the requirement for the crystalloid body, may manifest later in female development, and at a time point not well represented by *in vitro* culture. To test this, we conducted *in vivo* infection experiments in mice. Groups of three IFN*γ*KO mice were treated daily by gavage with 100 μl vehicle, or 10 mg/kg rapamycin, beginning two days prior to infection with 10,000 AP2-F-DiCre strain oocysts (Fig. 6A). Nluc activity was monitored in mouse feces, and we observed loss of oocyst shedding in mice treated with rapamycin compared to the vehicle control (Fig. 6B). We found this to be true in four independent biological experiments using groups of three mice each. Fig. 6C shows the area under the curve normalized for each matched group of treated and untreated mouse samples for all experiments (p<0.0001). Mice were euthanized on day six in two of these experiments, intestines were resected, and Nluc activity was measured for three ileal punch biopsies per mouse. In contrast to fecal shedding, intestinal Nluc readings were not reduced but were moderately higher in treated mice (Fig. 6D). To measure the effects of rapamycin treatment alone we used the TK-DiCre strain as a control (Fig. 6E-G, note that the TK gene is dispensable). We found rapamycin to be of no detriment to infection with these parasites, and luciferase readings were consistently higher upon treatment, regardless of whether we measured fecal or intestinal luciferase. This is likely due to the immunosuppressive effects of this drug (45). We conclude that AP2-F is critical to the formation and/or shedding of oocysts from infected animals, but that it is not required for the growth of parasites in the intestinal tissue.

**Figure 6:**
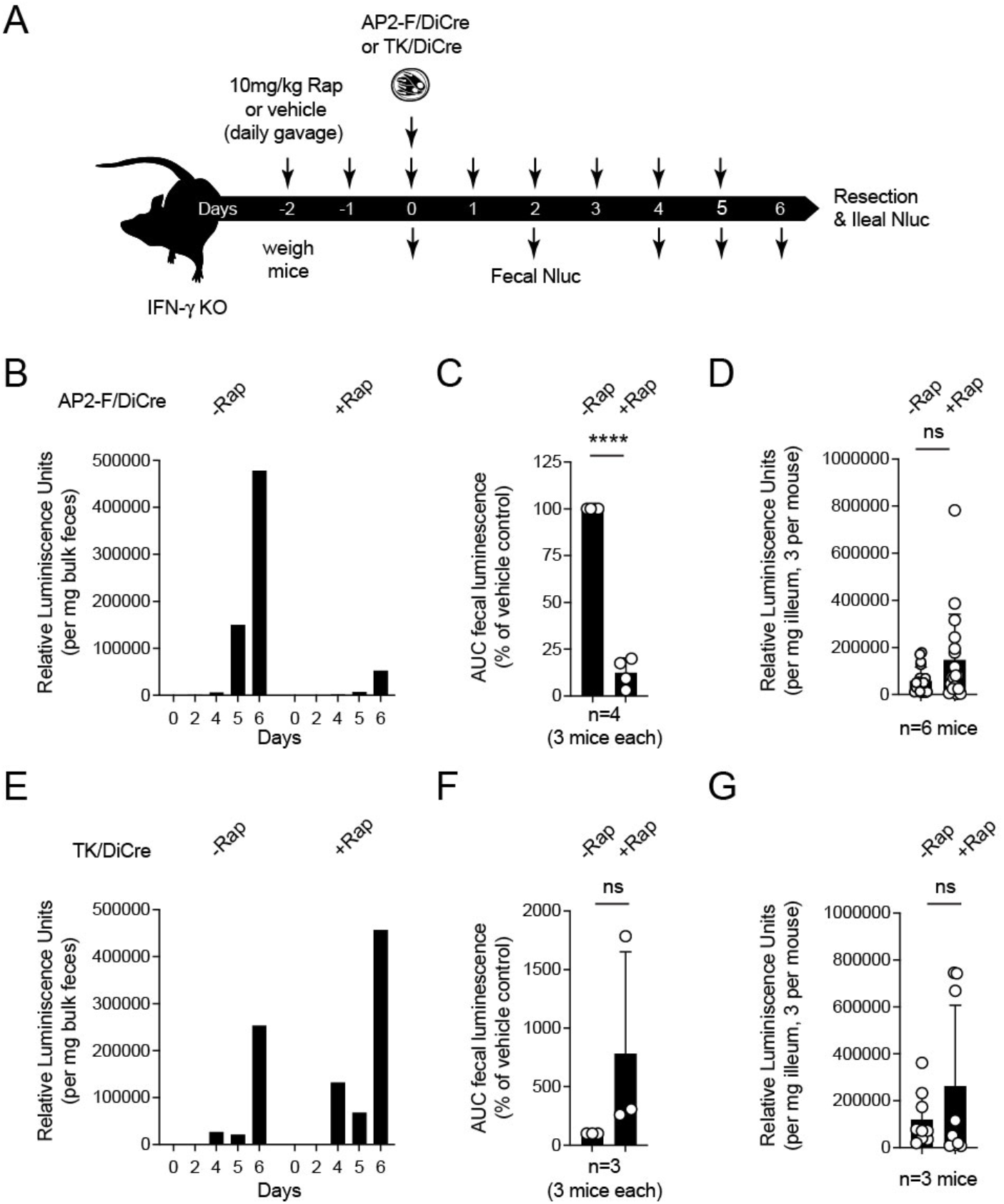
Ablation of AP2-F in infected animals leads to reduced oocyst shedding. (**A**) Schematic outline of treatment and sampling for infection of IFN-/-mice with AP2-F/DiCre (**B**-**D**) or TK/DiCre (**E**-**G**) parasites. (**B** and **E**) Luciferase measurements of feces collected from cages of three mice over the course of six days of infection. (**C** and **F**). Area under the curve for luciferase measurements as shown in (**B** and **E**) across 4 (**C**) and 2 (**F**) repeats. (**D** and **G**) Mice were euthanized on day 6 of infection, intestines were resected, flushed with buffer, and three ileal punch biopsies were taken per mouse and used in luciferase assays as a proxy for the intensity of intestinal infection. Symbols show individual mice or biopsies, respectively, and bars indicate mean and standard deviation. Significance was determined by unpaired student’s t-test.

## Discussion

New experimental tools have enabled a reevaluation of the *Cryptosporidium* lifecycle. The picture emerging from these studies, is that of a remarkably succinct developmental program, which relies on a single host, only three morphological stages, and unfolds over the course of only three days (2, 13, 15, 16). However, the molecular mechanisms by which this program is controlled remain poorly understood. In this study, we identified and characterized two AP2 transcription factors that are exclusively expressed in gametes. We demonstrated AP2-M to be a male factor, and not as initially assumed a female specific protein. Interestingly, labeling for the protein is narrowly restricted to the mitotic phase of male development, and lost prior to full maturation of the 16 male gametes. This is consistent with a model of the apicomplexan cell and developmental cycle driven by waves of gene expression, controlled by a succession of specific transcription factors (1, 24). Each phase is initiated by transcription and translation of the regulatory factor and terminated, at a defined point in the cell cycle, by degradation of that factor, thus yielding the field to the next transcription factor and the next wave of transcripts (46-48).

Genetic studies of developmental regulation in *Plasmodium* or *Toxoplasma* have been facilitated by the fact that transition to sex or cyst formation, respectively, is not essential to the growth of the lifecycle stages used to isolate mutants. In contrast, transgenesis in *Cryptosporidium* requires sporozoites, the products of sex (31). It is thus impractical to ablate transcription factors critical to lifecycle progression. To overcome this limitation, we adapted a DiCre conditional gene excision system that can be triggered by rapamycin treatment (40, 49), and used it to disrupt the AP2-F gene. Typically, such systems are assembled in multiple steps that first introduce the recombinase, and subsequently flank the target gene with loxP sites. The technical requirements of *C. parvum* transfection forced us to miniaturize the components into one cassette, to be delivered in a single genomic recombination event. Our strategy relied on short introns carrying LoxP sites, and we showed that those are well tolerated, and efficiently excised using two genes, TK, and AP2-F, as examples. To conditionally disrupt the large AP2-F gene, we replaced its promoter with a floxed promoter. Intergenic regions, including promoters are very short in *Cryptosporidium* (50), offering attractive targets for excision. Critical to implementing this strategy is to identify a surrogate promoter that will preserve the timing and strength of expression of the native gene. Multiple transcriptomic data sets facilitate this now (2, 32).

Consistent with previous reports in other systems (40, 49), we observed some ‘leaky’ DiCre activity, and we documented excision in the dispensable TK gene in the absence of rapamycin. We did not conduct rigorous serial passage experiments, but it is likely that the floxed segment is successively lost over time. In contrast, we did not observe such loss for the floxed AP2-F parasites in the absence of rapamycin induction. While there are several potential explanations for this difference, the most likely reason may be, that excision of an essential gene will result in loss of those parasites from the population, thus providing strong purifying counter-selection. When rapamycin was added to infected cultures, we observed inducible Cre activity in both strains and with similar kinetics. The number of parasites that express the AP2-F protein was reduced by 87% over 48h. Loss of AP2-F did not significantly affect the growth and asexual amplification of parasites in culture (or in mice), consistent with its role, being specifically restricted to the biology of the female gamete. Loss of the protein also did not impact commitment to female fate or development, as numbers of female gametes were indistinguishable between treated and control experiments. However, when we induced loss of AP2-F during infection in mice, we observed reduced shedding of oocysts that was not noted in a control strain. Overall, we conclude that AP2-F, and the genes under its control, are essential for the completion of the *C. parvum* lifecycle and required for the ability of the female gamete to undergo fertilization, or post-fertilization development leading to the formation and release of new infectious oocysts.

What may be the molecular nature of that step? Sequencing analyses of AP2-F in culture revealed reduced transcription of a focused set of genes, with a by-and-large exclusively female expression pattern. Among them were meiosis associated proteins including NIMA kinase. *Plasmodium berghei* homologs of this protein were shown to be required for meiosis and completion of the mosquito portion of the malaria life cycle (51, 52). However, most prominent among the transcripts lost, were those that encode most of the components of the crystalloid body. We recently defined the composition of this organelle, along with that of many other sporozoite compartments, in a proteomic experiment that used the hyperplexed localization of organelle proteins by isotope tagging (Hyper-LOPIT) approach (44, 53). We tested for gene set enrichment among the genes impacted by loss of AP2-F and found crystalloid body, but not oocyst wall proteins, to be highly enriched (Supplementary Fig. 5, p<0.001, FDR: 0). The crystalloid body is a multivesicular organelle containing an ordered array of spherical vesicles of 25-60 nm in diameter, and found at the basal end of *Cryptosporidium* sporozoites, but absent from merozoites (54, 55). Crystalloid bodies have been observed in other apicomplexans (55), and the structure is best characterized in *Plasmodium* species, where it is found in the post-fertilization ookinete and early oocyst stages (56). The molecular components of the crystalloid body are proteins that carry signal peptides, and feature complex interaction domains, including the CCp family of proteins which share similarity with the *Limulus* coagulation factor C, or Pleckstrin homology domain proteins, and several members have homologs in both *Plasmodium* and *Cryptosporidium* (57, 58). Importantly, mutants in these CB proteins, as well as related gametocyte proteins in *Plasmodium*, were demonstrated to suffer aberrant oocyst and/or sporozoite formation leading to loss of transmission from insect to mammal (57, 59). This is an intriguing parallel and consistent with the phenotype observed here, however, we currently lack antibody reagents in *C. parvum* to directly test such a role.

AP2 DNA binding proteins have been identified as regulators of stage differentiation, including female development and ookinete formation in *Plasmodium* (23, 24, 26, 60, 61), and we show in this study that they also control key steps in *Cryptosporidium* male and female biology. The simplicity of its single host lifecycle makes *C. parvum* a powerful model to dissect the fundamental biology of apicomplexan development. Conditional gene ablation as developed here, and induced protein destabilization as described in a recent study by Choudhary and colleagues (37), now open the door to rigorous genetic analysis of this fascinating process.

## Materials and Methods

### Mice

Ifng-/- mice (stock no:002287) were purchased from Jackson Laboratory and bred in house. Mice of both sexes were used (typically ranging from 6 to 8 weeks in age). Mice were treated with antibiotics, infected, and handled as detailed in (62).

### Cells

HCT-8 cells were purchased from ATCC (CCL-224TM) and maintained in RPMI-1640 medium (Sigma-Aldrich) supplemented with 10% Cosmic Calf Serum (HyClone), 1% penicillin-streptomycin (Gibco), 1% L-glutamine (Gibco), and 1x amphotericin B (Goldbio). For HCT-8 infection medium, serum concentrations were reduced to 1%.

### Reagents and drugs

If not indicated specifically reagents were purchased from Sigma-Aldrich. Rapamycin (LC Laboratories) was dissolved in 95% ethanol at a stock concentration of 50mg/ml. For *in vivo* studies, rapamycin was further diluted in water. Mice were weighed and treated daily by oral gavage with 100 μl of drug solution adjusted to deliver 10mg/kg (note that the ethanol concentration must not exceed 5% and was 2.5% in most of our experiments). A vehicle control was established from 95% ethanol by equivalent dilution.

### Antibodies and stains

Primary antibodies used in this study include rat anti-HA (Roche clone 3F10), VVL-FITC (1:1000, Vector FL1231), DMC1 antibody (1:10, a kind gift of Dr. Huston, University of Vermont (15)), rabbit anti-TrpB (31), and mouse anti-alpha tubulin (a kind gift of Jacek Gaertig, University of Georgia) were both used at 1:300, and secondary antibodies carrying a variety of Alexa Fluors (Abcam) at 1:1000.

### Plasmid construction

The DiCre cassette was built using a *P. falciparum* construct (40) kindly shared by Tobias Spielmann (Bernhard Nocht Insitut Hamburg, Germany). Guide oligonucleotides (Sigma-Aldrich) were introduced into the *C. parvum* Cas9/U6 plasmid16 by restriction cloning. See (63) for a detailed discussion of guide design for *C. parvum*.

Transfection plasmids were constructed by Gibson assembly using NEB Gibson Assembly Master Mix (New England Biolabs). Supplementary Table 1 provides the sequences of all primers used in this study.

### Parasite strains

We generated and purified transgenic parasite lines using *Cryptosporidium parvum* Iowa II strain oocysts (Bunchgrass Farms, Deary, ID) following protocols previously detailed (31, 62, 63).

### Nanoluciferase assay

The Nano-Glo® Luciferase Assay system (Promega) and Glomax 3000 (Promega) were used to measure luciferase activity in parasites, following the manufacturer’s protocol unless otherwise noted. For fecal samples, 19 – 21 mg of feces was processed as described previously in (63). For *in vitro* cultures, monolayers were resuspended in the NanoGlo substrate solution directly and luminescence measured. For intestinal samples, gut punches were weighed and lysed in fecal lysis buffer (63) by vortexing for 10 minutes with glass beads. Samples were mixed 1:1 with NanoGlo substrate solution and assayed in triplicate.

### Transient transfection assay for engineering DiCre conditional system

We followed the protocols for transient transfection of parasites as described (31). Briefly, 1 × 10^7^ *C. parvum* oocysts (Bunchgrass Farms, Deary, ID) were excysted and transfected with 10 ug each of DiCre and floxed plasmids using an Amaxa 4D Electroporator (Lonza). The transfected parasites were diluted in HCT-8 infection medium and divided equally to infect three 24 well plates of sub-confluent HCT-8 cells. Nanoluciferase activity was measured 48 hrs after infection.

### PCR assay for DiCre activation in stable transgenics

HCT-8 cells were infected with 50,000 oocysts in HCT-8 infection medium with and without 100 nM of rapamycin. After different times of infection, DNA was isolated from the cells according to the quick-start protocol from the DNeasy® Blood and Tissue Kit (Qiagen). PCR primers used are listed in Supplementary table 1.

### Immunofluorescence assay

HCT-8 cells were grown on coverslips in either 24- or 96-well plates and 20,000 of prepared oocysts in HCT-8 infection medium, respectively. For DiCre activation experiments, parasites were also grown in the presence of 100 nM rapamycin. Cells were fixed with 4% paraformaldehyde (Electron Microscopy Science) in PBS and then permeabilized with PBS containing 0.25% Triton X-100. The cells were blocked with 3% bovine serum albumin (BSA, Sigma-Aldrich) solution, followed by incubation with primary antibodies. Cells were washed with PBS and then incubated with appropriate fluorophore-conjugated secondary antibodies and counterstained with DAPI (Thermo-Fisher). Coverslips were then mounted on glass slides with fluorogel (Electron Microscopy Science) mounting medium. Super-resolution structured illumination microscopy was conducted using a Carl Zeiss Elyra (UGA Biomedical Microscopy Core) or a GE OMX (PennVet Imaging Core) microscope. Widefield microscopy was performed using a Leica LAS X microscope (PennVet Imaging Core) and images were processed and analyzed using Carl Zeiss ZEN v.2.3 SP1, GE Softworx, and NIH ImageJ software.

### EdU labelling to measure thymidine kinase activity

HCT-8 cells were infected with 100,000 oocysts of the TK Flox Dicre strain. EdU (Thermo-Fisher) was added to cultures 36 h after infection to a final concentration of 10 μM and cells were fixed 12 h later. A Click-iT EdU Alexa-Fluor 594 kit (Thermo Fisher Scientific) was used to label incorporated EdU. Parasites were counter-stained with anti-HA antibody and fluorescein conjugated *Vicia villosa* lectin (Vector Laboratories).

### Rapamycin induction *in vivo*

Four-to-six week-old female IFNγ^−^/^−^ mice were treated with 10mg/kg of rapamycin (LC Laboratories) via oral gavage daily, starting 2 days prior to infection, and continuing 5 days post infection. Mice were sex and age matched between treatment groups. Following two days of pre-treatment with rapamycin, mice were gavaged with 10,000 oocysts in PBS. Bulk fecal samples were collected, and intestinal punches were harvested on day 6. Punches were stored at −80 ºC prior to processing for nanoluciferase measurement.

### RNA sequencing

HCT-8 cells were infected with 100,000 oocysts of wild type (non-transgenic) *C. parvum* or AP2-F DiCre strains with or without 100nM rapamycin. Cells were lysed and RNA extracted using the RNeasy Mini Kit (Qiagen, Hilden, Germany) according to the manufacturer’s protocol. Illumina Stranded mRNA Prep, Ligation kit (20040534) was used for cDNA generation and library preparation. Total RNA and libraries were quality controlled and quantified using the Tapestation 4200 (Agilent Technologies, Santa Clara, CA) and Qubit 3 (Thermo Fischer Scientific, Waltham, MA) instruments. Samples were pooled and single-end reads were generated on an Illumina NextSeq 500 sequencer.

### RNA-sequencing analysis

Raw reads were mapped to the *C. parvum* Iowa II reference (VEuPathDB, release 50) using Kallisto v.0.45.0 (64). All subsequent analyses were carried out using the statistical computing environment R v.4.0.3 in RStudio v.1.3.1093 and Bioconductor. In brief, transcript-level quantification data were summarized to genes using the tximport package and data were normalized using the TMM method, which was implemented in EdgeR (65). Only genes with more than 10 counts per million in at least 3 samples were used for analysis. Precision weights were applied to each gene based on the mean–variance relationship using the VOOM function in Limma (66). Linear modelling and Bayesian statistics carried out in Limma were used to identify differentially expressed genes with a FDR-adjusted p-value of ≤0.01 and an absolute log2-transformed fold change of ≥1 after adjustment with Benjamini– Hochberg correction. Genes were identified as oocyst wall and crystalloid body based on analyses derived from the *C. parvum* Hyper-LOPIT proteomic dataset (44). Gene set enrichment analysis was carried out using GSEA software 4.3.2 (67). Oocyst wall and crystalloid body signatures obtained from the Hyper-LOPIT dataset were used for GSEA with 1000 permutations, a weighted enrichment statistic, and the “Diff_of_Classes” metric for ranking genes. All other parameters were kept as the default settings.

### Statistical method

GraphPad PRISM was used for all statistical analyses. When measuring the difference between the two populations, we used a standard Student’s t-test. No statistical tests were used to predetermine sample size and no animals were excluded from results.

ANOVA was used to compare the means of multiple groups followed by Tukey’s post hoc test for pair-wise comparisons.

### Animal ethics statement

All the protocols for animal experimentation were approved by the Institutional Animal Care and Use Committee of the University of Georgia (protocol A2016 01-028-Y1-A4) and/or the Institutional Animal Care and Use Committee of the University of Pennsylvania (protocol number 806292). No statistical tests were used to predetermine the sample size of mice used for experiments. Mice were not randomized, and investigators were not blinded before any of the experiments.

## Data and code availability

RNA-sequencing data generated in this study are available from GEO database repository under accession number GSE216844. All code used in these analyses is available in Supplementary File 2 and on GitHub (https://github.com/katelyn-walzer/Ablation_of_AP2-F_Manuscript).

## Acknowledgements

This work was supported in part by grants from the National Institutes of Health to BS (R01AI127798 and R01AI112427) and a postdoctoral fellowship to KAW (F32 AI154666). We thank Tobias Spielmann for DiCre plasmids and advice, Chris Huston and Jacek Gaertig for antibodies, and Elise Krespan from CHMI for help with library preparation and sequencing. We thank the UGA Biomedical Microscopy Core and the Penn Vet Imaging Core for support of microscopy, and Carrie Brooks, Emily Kugler, Briana McLeod and Eleanor Smith for help with in vivo experiments.

---

**Supplementary Table 1:**
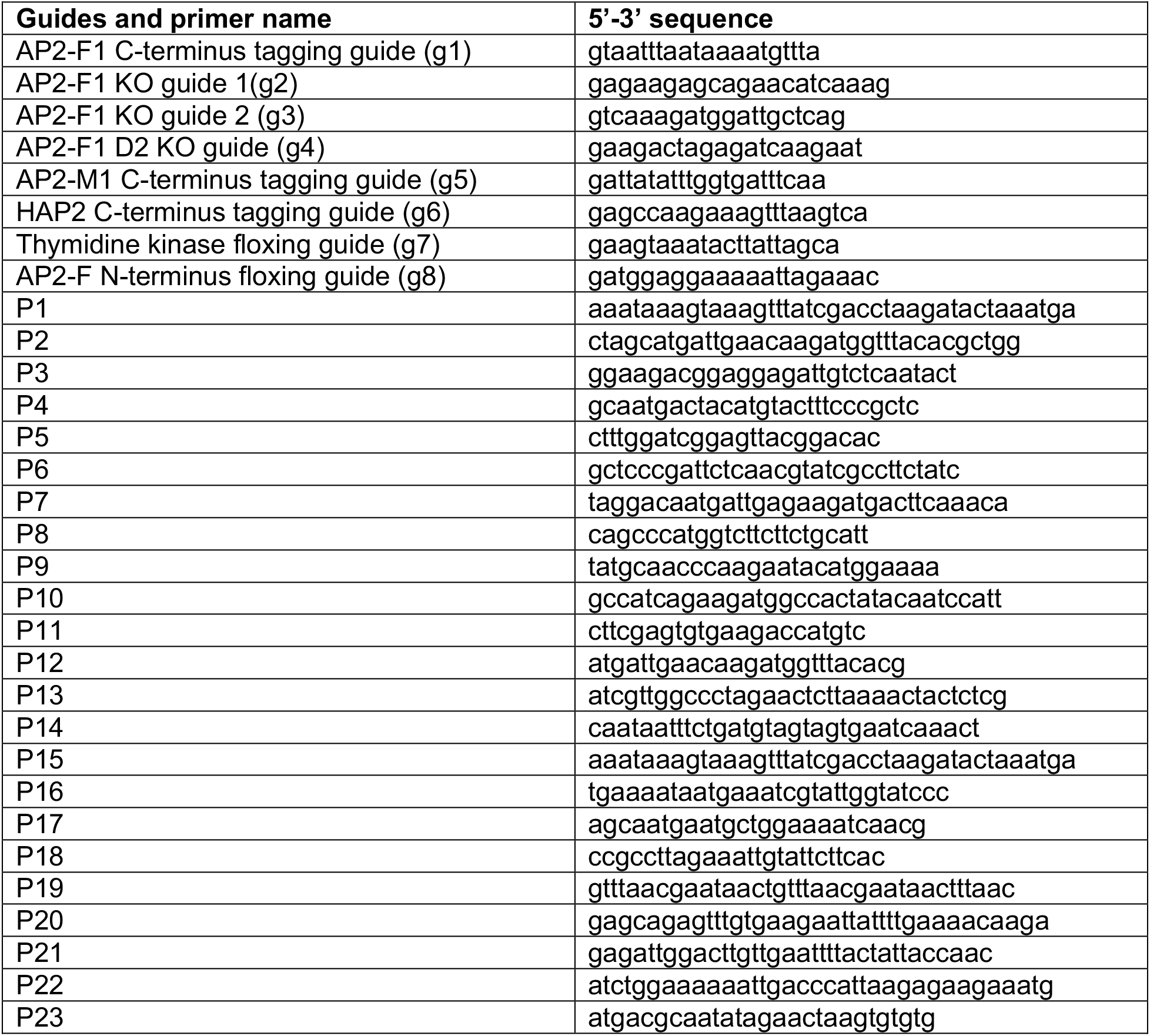
Primers used in this study.

**Supplementary Fig 1:**
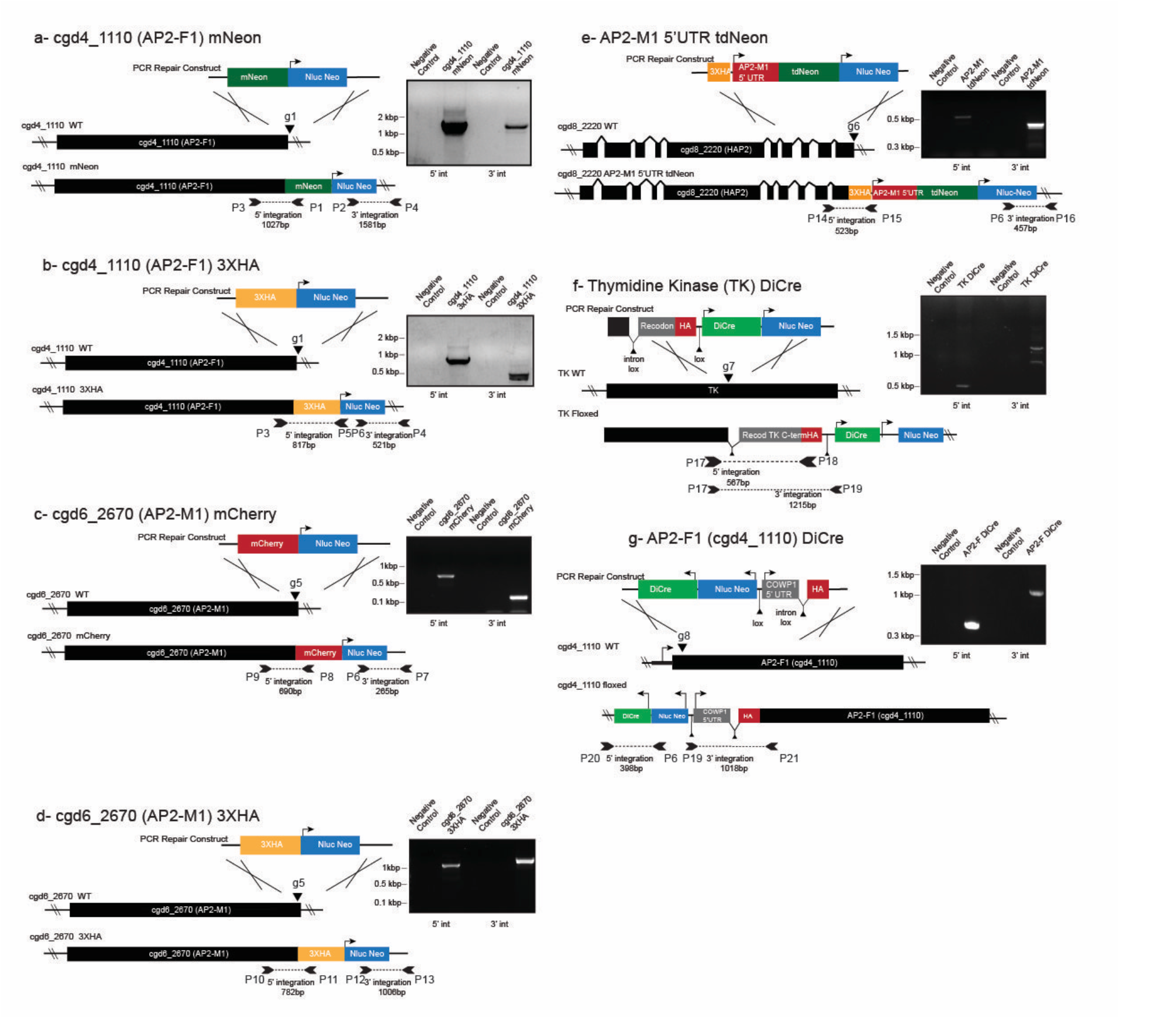
Integration maps and validation PCR of all strains engineered. Diagnostic PCR products are shown as inverted arrows. Note that maps are not to scale and that primers indicated here are listed in supplementary table 1.

**Supplementary Fig 2:**
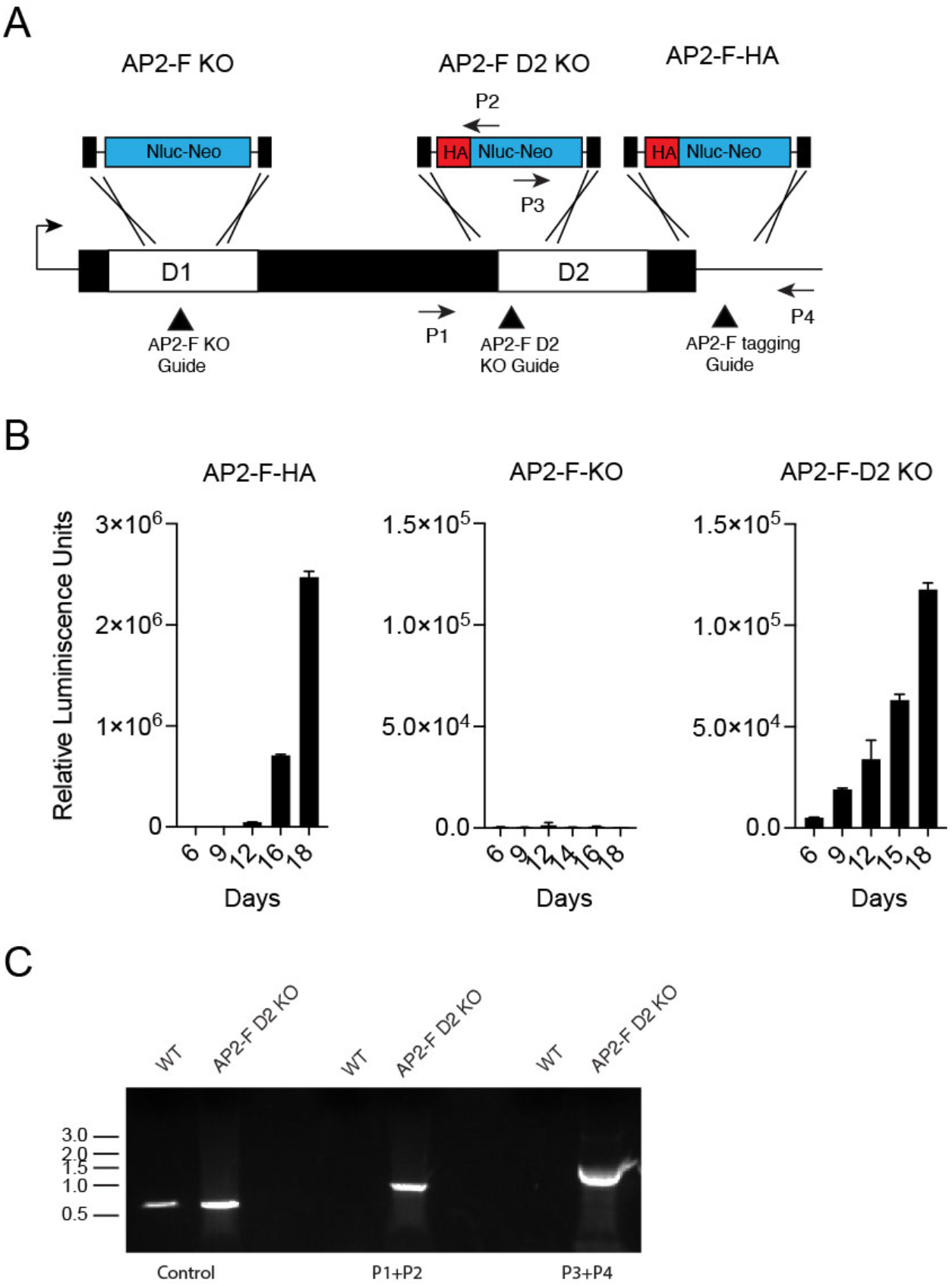
The AP2-F locus is refractory to disruption of the N-terminal AP2 DNA binding site. (**A**) Map showing three different modification of the AP2-F locus that were attempted here: deletion of the N-terminal or C-terminal AP2 DNA binding domain or C-terminal epitope tagging. (**B**) Luciferase measurements of feces collected from mice infected sporozoites transfected with the indicated vectors. Note that while viable parasites emerged from selection for AP2-F epitope tagging and KO of the C-terminal domain, that was not the case for the N-terminal deletion (which ablates essentially the entire gene). (**C**) PCR analysis demonstrating successful insertion into the portion of the AP2-F gene encoding the C-terminal AP2 DNA binging domain.

**Supplementary Fig 3:**
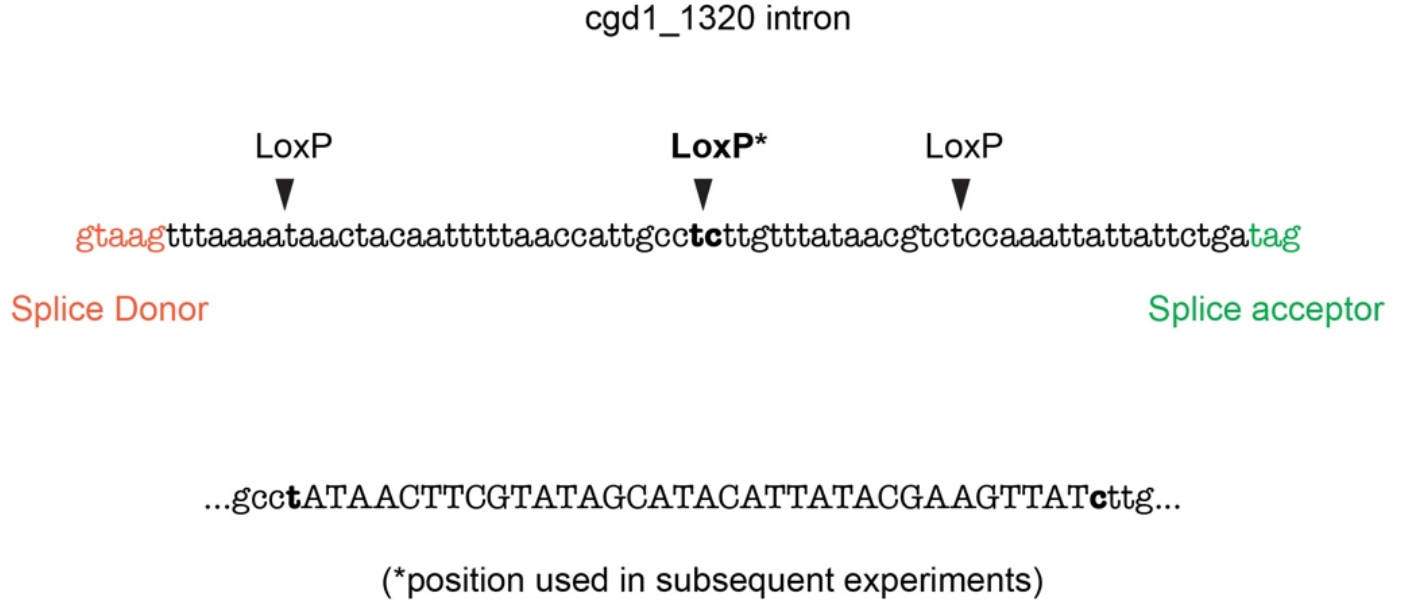
The floxed artificial intron used in this study. The position of three loxP sites within the intron found in the *C. parvum* gene cgd1_1320 that tolerate insertion of a LoxP site are shown. The LoxP site shown in bold was used for subsequent constructs, and the insert below shows the LoxP (upper case) insertion in detail.

**Supplementary Fig 4:**
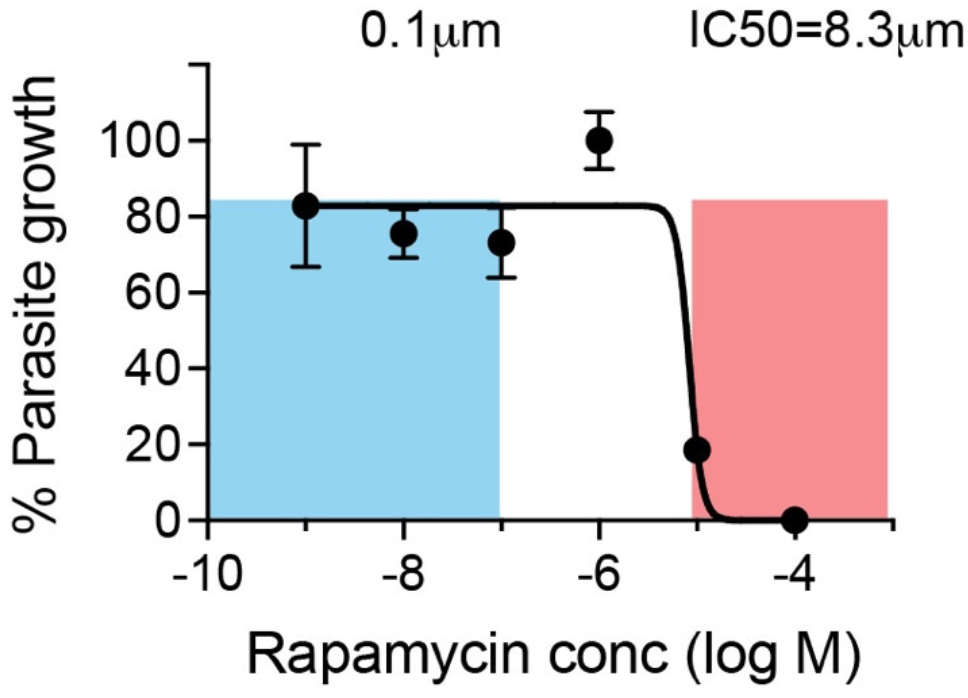
Impact of rapamycin for *Cryptosporidium* growth in culture. HCT-8 cell cultures were infected with *C. parvum* expressing Nluc luciferase and grown in the presence of different concentrations of rapamycin for 48h. Growth was measured by luciferase assay (each symbol shows the mean of three biological replicates and the error bar shows the standard deviation, values were normalized with the highest measurement set to 100%). The IC50 was determined using Prism 9 software and a non-linear fit model shown as a line here. Concentrations above the IC50 are highlighted in red. Concentrations used in this study (100 nM) and lower are highlighted in blue and we note that in the range used rapamycin does not impact parasite growth.

**Supplementary Fig 5:**
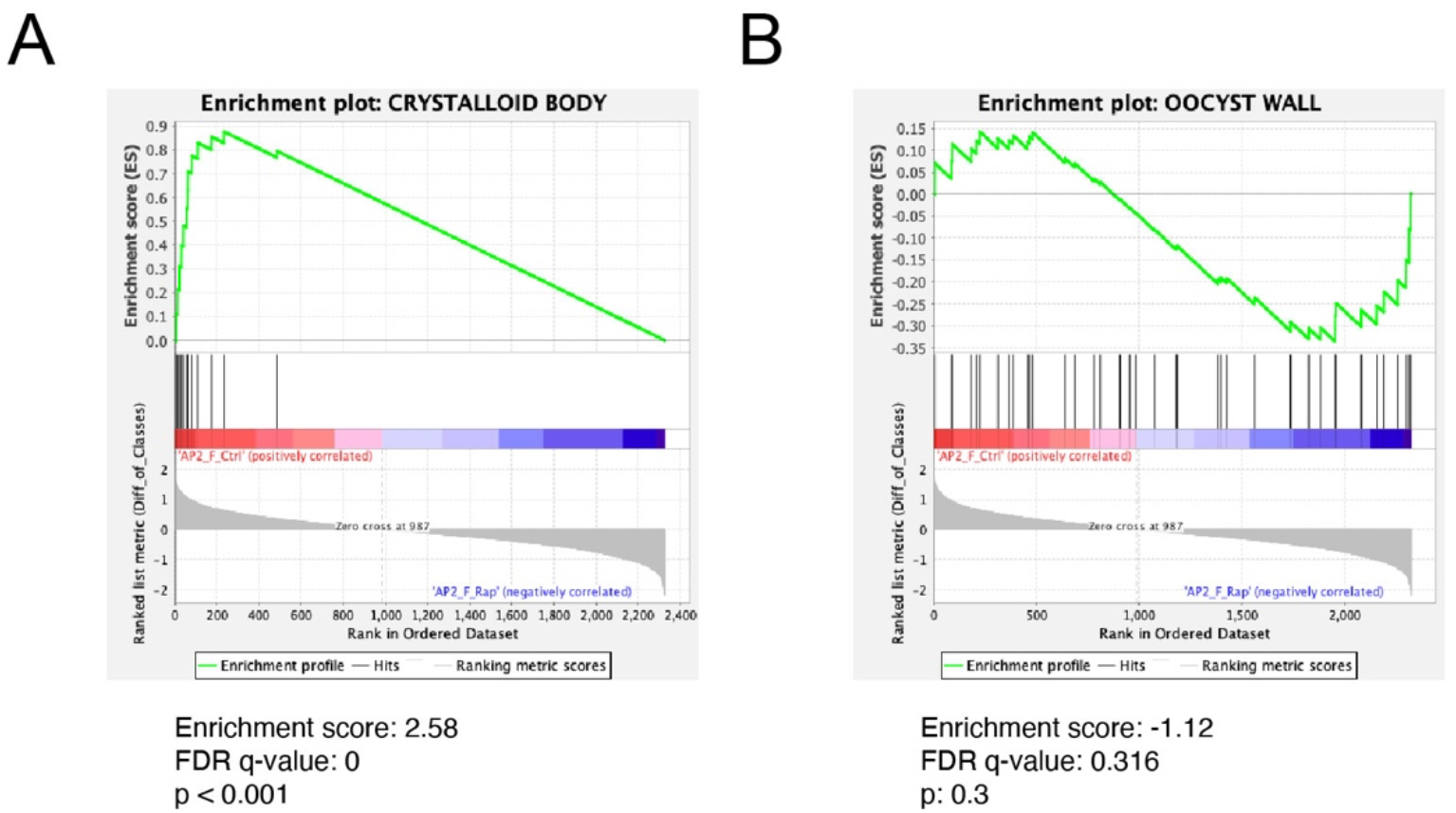
Gene set enrichment analyses of female gene signatures upon the loss of AP2-F. Analyses were conducted using GSEA 4.3.2 to show enrichment in the untreated control (reflecting loss upon rapamycin treatment). Plots shown use the Hyper-LOPIT defined (44) *C. parvum* gene sets for the crystalloid body (A) and oocyst wall (B, as a female specific control).

**Supplementary File 1**. Differentially Expressed Genes. Multi-tab Excel file containing differentially expressed genes between all samples. First tab contains all differentially expressed genes and other tabs contain comparisons between individual samples. AP2-F is highlighted in red. Crystalloid body proteins are highlighted in magenta. Expression data is normalized counts per million and log2 transformed.

**Supplementary File 2**. Supplementary code file.

